# *Clostridium difficile* colonizes alternative nutrient niches during infection across distinct murine gut microbiomes

**DOI:** 10.1101/092304

**Authors:** Matthew L. Jenior, Jhansi L. Leslie, Vincent B. Young, Patrick D. Schloss

## Abstract

*Clostridium difficile* is the largest single cause of hospital-acquired infection in the United States. A major risk factor for *Clostridium difficile* infection (CDI) is prior exposure to antibiotics, as they disrupt the gut bacterial community which protects from *C. difficile* colonization. Multiple antibiotic classes have been associated with CDI susceptibility; many leading to distinct community structures stemming from variation in bacterial targets of action. These community structures present separate metabolic challenges to *C. difficile.* Therefore we hypothesized that the pathogen adapts its physiology to the nutrients within different gut environments. Utilizing an *in vivo* CDI model, we demonstrated *C. difficile* highly colonized ceca of mice pretreated with any of three antibiotics from distinct classes. Levels of *C. difficile* spore formation and toxin activity varied between animals based on the antibiotic pretreatment. These physiologic processes in *C. difficile* are partially regulated by environmental nutrient concentrations. To investigate metabolic responses of the bacterium *in vivo*, we performed transcriptomic analysis of *C. difficile* from ceca of infected mice across pretreatments. This revealed heterogeneous expression in numerous catabolic pathways for diverse growth substrates. To assess which resources *C. difficile* exploited, we developed a genome-scale metabolic model with a transcriptome-enabled metabolite scoring algorithm integrating network architecture. This platform identified nutrients *C. difficile* used preferentially between pretreatments, which were validated through untargeted mass spectrometry of each microbiome. Our results supported the hypothesis that *C. difficile* inhabits alternative nutrient niches across cecal microbiomes with increased preference for nitrogen-containing carbon sources, particularly Stickland fermentation substrates and host-derived glycans.

**Importance:** Infection by the bacterium *Clostridium difficile* causes an inflammatory diarrheal disease which can become life-threatening, and has grown to be the most prevalent nosocomial infection. Susceptibility to *C. difficile* infection is strongly associated with previous antibiotic treatment, which disrupts the gut microbiota and reduces its ability to prevent colonization. In this study we demonstrated that *C. difficile* altered pathogenesis between hosts pretreated with antibiotics from separate classes, and exploited different nutrient sources across these environments. Our metabolite score calculation also provides a platform to study nutrient requirements of pathogens during an infection. Our results suggest that *C. difficile* colonization resistance is mediated by multiple groups of bacteria competing for several subsets of nutrients and could explain why total reintroduction of competitors through fecal microbial transplant currently is the most effective treatment for recurrent CDI. This work could ultimately contribute to the identification of targeted, context-dependent measures that prevent or reduce *C. difficile* colonization including pre- and probiotic therapies.

## Introduction

Infection by the Gram-positive, spore-forming bacterium *Clostridium difficile* has increased in both prevalence and severity across numerous countries during the last decade (1). In the United States, *C. difficile* was estimated to have caused >500,000 infections and resulted in ∼$4.8 billion worth of acute care costs in 2014 (2). *C. difficile* infection (CDI) causes an array of toxin-mediated symptoms ranging from abdominal pain and diarrhea to the more severe conditions pseudomembraneous colitis and toxic megacolon. Prior treatment with antibiotics is the most common risk factor associated with development of CDI (3). Antibiotics contribute to an individual’s susceptibility to CDI by disrupting their gut microbiota (4). In mouse models, multiple antibiotics can induce susceptibility to *C. difficile* colonization (5–7). Notably, each antibiotic resulted in unique gut bacterial communities that permitted to high levels of *C. difficile* colonization. Others have also shown that antibiotics from multiple classes also alter the gut metabolome, increasing the concentrations of some *C. difficile* growth substrates (6, 8–10). The ability of an unaltered murine gut community to exclude *C. difficile* colonization supports the nutrient-niche hypothesis, which states that an organism must be able to utilize a subset of available resources better than all competitors to colonize the intestine (11, 12). Taken together these results are a strong indication that the healthy gut microbiota inhibits the growth of *C. difficile* by limiting the availability of the substrates it needs to grow.

Based on genomic and *in vitro* growth characteristics, *C. difficile* appears able to adapt to a variety nutrient niches (13). *C. difficile* has a relatively large and mosaic genome, it can utilize a variety of growth substrates, and possesses a diverse array host range (6, 14–16). These qualities are hallmarks of ecological generalists (17). *C. difficile* has also been shown to integrate signals from multiple forms of carbon metabolism to regulate its pathogenesis. *In vitro* transcriptomic analyses suggest that high concentrations of easily metabolized carbon sources, such as glucose or amino acids, inhibit toxin gene expression and sporulation (18, 19). Other studies have indicated that other aspects of *C. difficile* metabolism may be influenced through environmental nutrient concentration-sensitive global transcriptional regulators such as CodY and CcpA (20, 21). These analyses focused on *in vitro* growth (22, 23) or colonization of germ-free mice (14, 21). Although these analyses are informative, they are either directed toward the expression of pathogenicity factors or lack the context of the gut microbiota which *C. difficile* must compete against for substrates. Metabolomic investigations have also been used to assay changes in bacterial metabolism as they relate to CDI and have characterized the levels of germinants and growth substrates (6, 10); however, metabolomic approaches are unable to attribute a metabolite to specific organisms in the gut community. Thus metabolomics more closely represents the echoes of total community metabolism, not the currently active processes of any one population. It has thus far not been possible to study *C. difficile*’s metabolism *in vivo*. To overcome these limitations, we implemented transcriptomic and untargeted metabolomic analyses of *C. difficile* and the surrounding environment to better understand the active metabolic pathways in a model of infection. Based on the ability of *C. difficile* to grow on a diverse array of carbon sources and its ability to colonize a variety of communities, we hypothesized that *C. difficile* adapts its metabolism to fit the context of the environment it is attempting colonize. To test this hypothesis, we employed a mouse model of infection to compare the metabolic response of *C. difficile* to the gut environment caused by three antibiotics from distinct classes. By characterizing a transcriptome-enabled metabolic model of *C. difficile* and changes in the metabolome of each respective environment, we were able to generate a systems model to directly test the nutrient-niche hypothesis.

## Results

**Levels of *C. difficile* sporulation and toxin activity vary among different microbiomes.** Conventionally-reared SPF mice were treated with either streptomycin, cefoperazone, or clindamycin (Table 1 and Fig. S1). These antibiotics were selected because they each have distinct and significant impacts on the structure of the cecal microbiome (Fig. S2A and S2B). We challenged the antibiotic treated mice and germ-free (exGF) mice with *C. difficile* stain 630 to understand the pathogen’s physiology with and without other microbiota. This toxigenic strain of *C. difficile* was chosen for its moderate clinical severity in mouse models (24) and well-annotated genome (25). After infection, we measured sporulation and toxin production at 18 hours post inoculation. That time point corresponded with when another laboratory strain of *C. difficile* reached its maximum vegetative cell density in the cecum with limited sporulation (26). There was not a significant difference in the number of vegetative *C. difficile* cells in the ceca of mice pretreated with any of the three antibiotics (Fig. 1A). All antibiotic treated and exGF mice were colonized to ∼1×10^8^ colony forming units (cfu) per gram of cecal content, while untreated mice maintained colonization resistance to *C. difficile* (Fig. 1A). Despite having the same number of vegetative *C. difficile* cells, more spores were detected in exGF mice than in the antibiotic pretreated mice (*P* = 0.003, 0.004, and 0.003; Fig. 1B). Toxin activity was relatively low across each group tested compared to previous studies (24, 27). The low activity was likely the result of the early sampling time point during infection. In spite of this, the toxin activity in exGF animals was significantly higher than any other colonized group (all *P* < 0.001), with slight variation between antibiotic pretreatment groups (Fig. 1C). These results showed that *C. difficile* colonized different communities to consistently high levels, but had subtle variation in sporulation and toxin activity between distinct antibiotic-pretreated environments. As activation of both traits has been linked to recognition of distinct nutrient source concentrations in the environment (28, 29), we hypothesized that *C. difficile* was utilizing different growth substrates across the conditions tested. To investigate the physiology of *C. difficile* when colonizing distinct susceptible gut environments, we performed whole transcriptome analysis of *C. difficile* from the cecal content of the same mice used to measure *C. difficile* density and toxin activity.

**Table 1.** **Antibiotics used during *C. difficile* murine infection models.**

| Antibiotic | Class | Target | Activity | Administration | Dosage |
| --- | --- | --- | --- | --- | --- |
| Cefoperazone | Cephalosporin (3rd generation) | Primarily Gram-positive bacteria, with increased activity against Gram-negative bacteria | Irreversibly crosslink bacterial transpeptidases to peptidoglycan and prevents cell wall synthesis | Drinking water Ad libitum for 5 days, 2 days untreated drinking water prior to infection | 0.5 mg/ml drinking water |
| Streptomycin | Aminoglycoside | Active against most Gram-negative aerobic and facultative anaerobic bacilli | Protein synthesis inhibitor through binding the 30S portion of the 70S ribosomal subunit | Drinking water Ad libitum for 5 days, 2 days untreated drinking water prior to infection | 5.0 mg/ml drinking water |
| Clindamycin | Lincosamide | Primarily active against Gram-positive bacteria, most anaerobic bacteria, and some mycoplasma | Protein synthesis inhibition through binding to the 23s portion of the 50S ribosomal subunit | Intraperitoneal injection 24 hours prior to infection | 10 mg/kg body weight |

**Figure 1.**
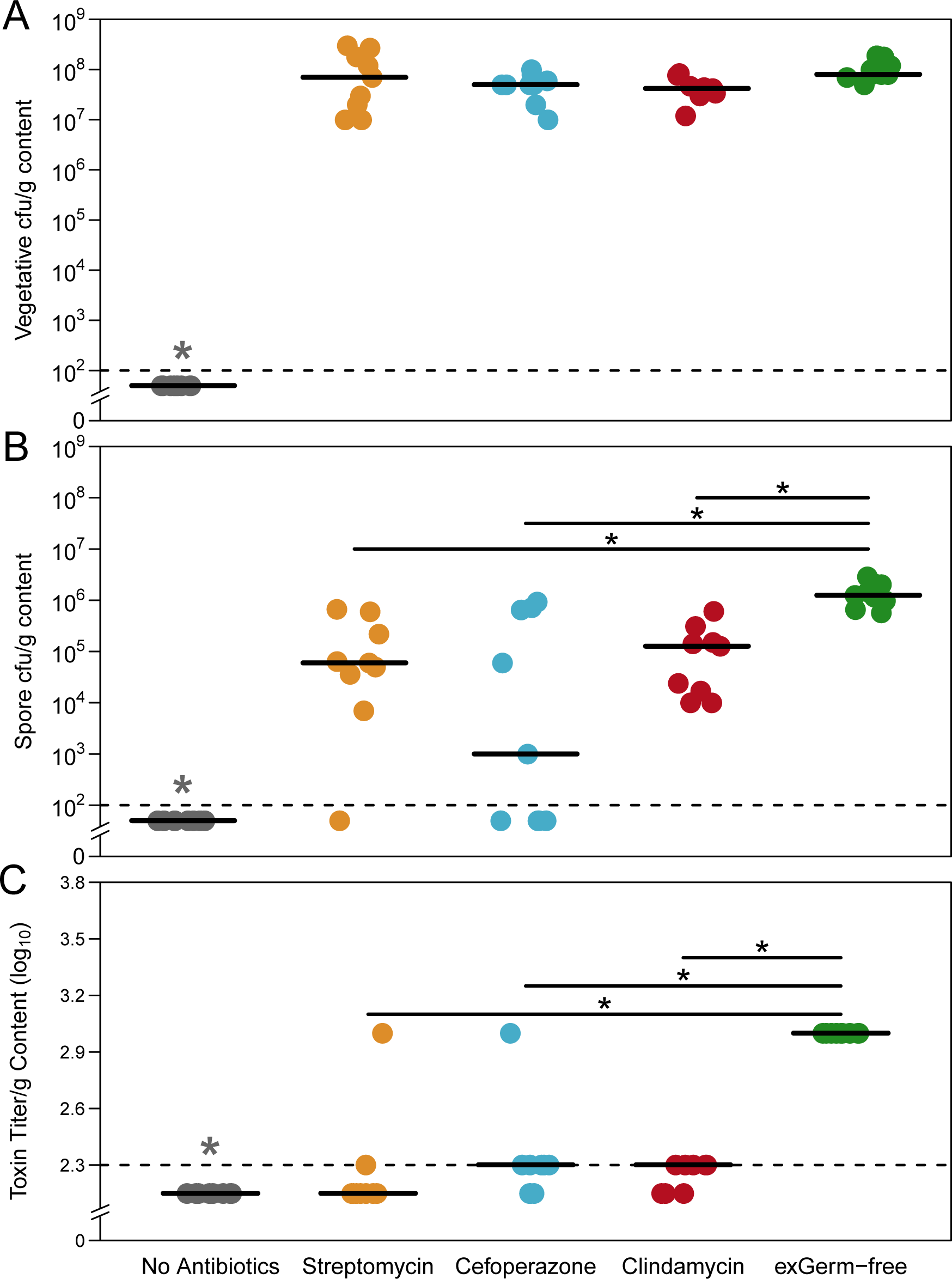
Gut environment context affects *C. difficile* sporulation and toxin activity. Quantification of spore cfu and toxin titer from cecal content of infected mice (n = 9 per group). **(A)** Vegetative *C. difficile* cfu per gram of cecal content (*P* = n.s.). **(B)** *C. difficile* spore cfu per gram of cecal content. **(C)** Toxin titer from cecal content measured by activity in Vero cell rounding assay. Dotted lines denote limits of detection (LOD). Values for undetectable points were imputed as half the LOD for calculation of significant differences. Significance (*P* < 0.05), denoted by single asterisk, was determined with Wilcoxon signed-rank test with Benjamini–Hochberg correction.

***C. difficile* alters its gene expression pathways when colonizing distinct antibiotic-pretreated environments.** Utilizing aliquots of cecal content from the same mice that were in the previous assays, we measured differential expression of specific genes associated with *in vivo* phenotype changes reported in previous studies with an RNA-Seq-based approach. Microarray-based gene expression measurement was not a viable alternative to sequencing as the amount of background orthologous transcription from other bacterial species would contribute to non-specific binding and bias the true *C. difficile* signal. Therefore we employed RNA-Seq to quantify *C. difficile*-specific transcription. *C. difficile* represented a small percentage of the community in each colonized environment (Fig. S2C), which made it impossible to sequence the transcriptome of individual mice due to the depth required to sufficiently sample the transcripts of *C. difficile*. This required the generation of a single transcriptome per condition using pooled mRNA from all mice within each pretreatment group. Following sequencing, read curation, and stringent mapping to *C. difficile* str. 630 genes (Materials & Methods) we implemented two steps of abundance normalization to compare expression between groups. Transcript abundances for each target gene were first corrected to both read length and target gene length, which resulted in an average per-base expression level for each. Adjusted values were then down-sampled to the same total read abundance for each mapping effort, allowing for even comparison between the conditions. Additionally, before proceeding with the analysis we did and assessed variation in expression of select bacterial housekeeping genes across treatment groups (Fig. S3A). Due to the heterogeneity of *C. difficile* reference genes across strains (30), we chose DNA gyrase subunit A (GyrA), threonyl-tRNA synthetase (ThrS), and ATP-dependent Clp protease (ClpP) because they are conserved across bacterial phyla and have been commonly utilized as standards for numerous transcriptional studies (14, 31, 32). We observed consistent expression for each of the housekeeping genes was observed across treatments, which indicated that our results were more likely to be a true reflection of *C. difficile* expression *in vivo*. We then focused on select genes previously demonstrated to have altered transcription based on environmental cues including several key sigma factors (29) and downstream genes involved in sporulation (33), toxin production (34), and quorum sensing (35) (Fig. S4). Comparing these data to results from the previous section, toxin gene expression seemed to vary between conditions more than the activity data would suggest (Fig. S4B). However, the relative abundance of cDNA transcript abundance recruited within this mapping effort to the toxin genes was very low, which would agree with the generally low levels of toxin activity detectable across treatment groups (Fig. 1C). For the other gene categories, consistent trends across pretreatments were not apparent through this analysis so we decided to shift our focus toward differences in metabolic pathways that were more explicitly involved in the breakdown of environmentally acquired nutrients.

We chose to assess transcriptional differences in several specific families of genes known to contribute to different aspects of *C. difficile* metabolism (Fig. 2A & Table S1). Genes involved in amino acid catabolism, including those that encoded enzymes involved in Stickland fermentation and general peptidases, had the highest level of expression across all pretreatments. Stickland fermentation refers to the coupled fermentation of amino acid pairs in which one is deaminated and the other is reduced to ultimately generate ATP (36). This suggested that *C. difficile* catabolized environmental amino acids during infection, regardless of the structure of the surrounding community. Although there were gene categories that were equally expressed across conditions in spite of the community differences, there were patterns of expression for certain gene families and specific genes that were distinct to each antibiotic pretreatment. In mice pretreated with cefoperazone, *C. difficile* tended to have more expression of genes in the ABC sugar transporter and sugar alcohol catabolism (e.g. mannitol) families and fewer genes in the PTS transporter family than the other pretreatment groups. In mice pretreated with clindamycin, *C. difficile* tended to have higher expression of genes from disaccharide catabolism (e.g. beta-galactosidases and trehalose/maltose/cellibiose hydrolases), fermentation product metabolism (including consumption or production of acetate, lactate, butyrate, succinate, ethanol, and butanol), and PTS transporter families. Genes from the sugar alcohol catabolism and ABC sugar transporter families were not highly expressed in the clindamycin-pretreated mice. Finally, in mice pretreated with streptomycin, *C. difficile* had higher levels of expression of genes from the sugar alcohol catabolism (e.g. sorbitol) and PTS transporter families. Combined, these results suggested that while catabolism of amino acids and specific carbohydrates are core components of the *C. difficile* nutritional strategy during infection, *C. difficile* adapted its metabolism across different susceptible environments.

**Figure 2.**
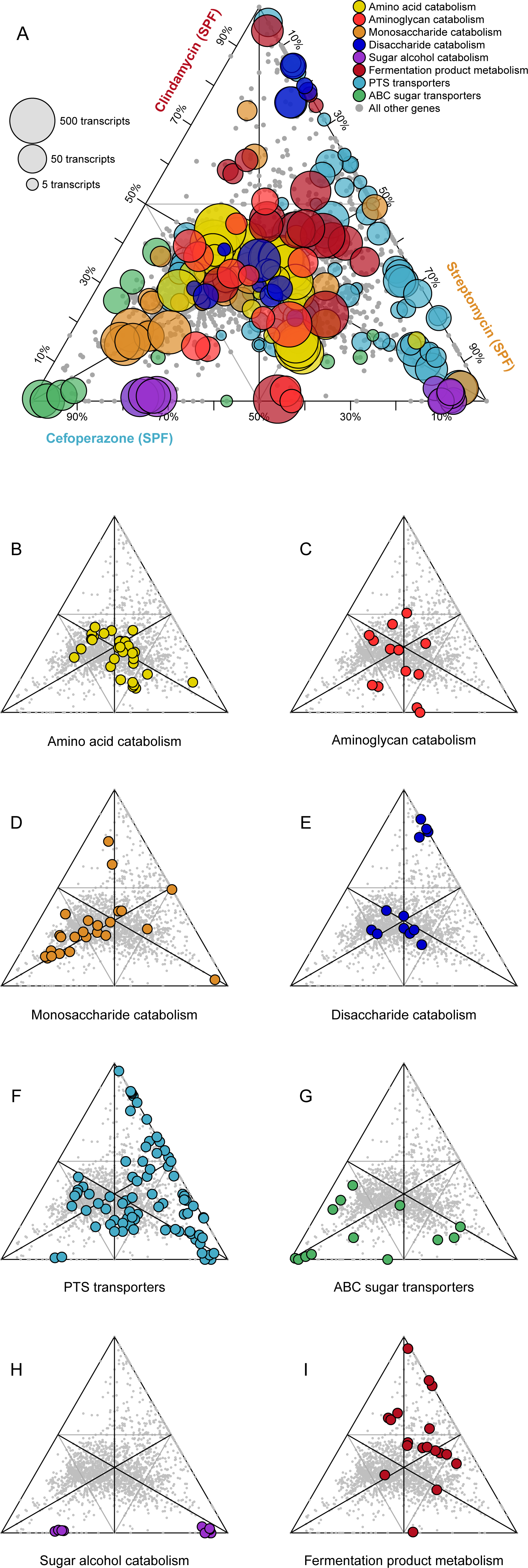
*C. difficile* alters expression metabolic pathways between antibiotic pretreatment models. Each point in the ternary plot represents a unique gene from the annotated genome of *C. difficile* str. 630. Position reflects the ratio of median rarefied transcript abundance for that gene between the three colonized antibiotic pretreatment models. Genes from specific metabolic pathways of interest are labeled and transcription from all other genes are shown in gray. **(A)** Size of highlighted points is relative to the largest transcript abundance among the antibiotic pretreatments for each gene. Categories of metabolism are displayed separately in **(B-I)**. Genes, annotations, and normalized transcript abundances can be found in Table S1.

**Genome-scale metabolic model structure underscores known *C. difficile* physiology.** Because multiple enzymes can utilize the same input substrates within a single organism, we decided to implement a metabolic network-based approach to further investigate which metabolites were differentially utilized between conditions by *C. difficile*. This approach is more robust at identifying reporter metabolites than assessing individual gene transcription because if the amount of a single enzyme that acts on a substrate decreases, yet others that also act on that substrate increase, those changes are more readily apparent in the context of a network. To perform this analysis, we created a generalizeable tool to generate *de novo* genome-enabled bipartite metabolic models with directed enzymatic reactions of bacterial species using KEGG gene and biochemical reaction annotations. We implemented this platform using the genome of *C. difficile* str. 630 (Fig. 3A), with enzymes and metabolites represented by nodes and their interactions by directed connecting edges. The *C. difficile* str. 630 network contained a total of 447 enzymes and 758 metabolites, with 2135 directed edges (Fig. 3A). To validate our metabolic network, we analyzed network topology by calculating two metrics of centrality, betweenness centrality (BC) and closeness centrality (CC), to determine which nodes are critical to the structure of the metabolic network and if these patterns reflect known biology (Table S3). Both metrics utilize shortest paths, which refer to fewest possible number of network connections that lie between two given nodes. The BC of each node is the fraction of shortest paths that pass through that node and connect all other potential pairs of nodes. In biological terms, this refers to the amount of influence a given hub has on the overall flow of metabolism (37). Similarly, CC is the reciprocal sum of the lengths of shortest paths included in each node’s BC. This value demonstrates how essential a given node is to the overall structure of the metabolic network (38). Metabolic network structural studies of *Escherichia coli* have found that metabolites with the highest centrality calculations are involved in fundamental processes in metabolism, namely glycolysis and the citrate acid cycle pathway (39). As such, these metrics allow for assessment of the degree to which a metabolic network accurately depicts established principles of bacterial metabolism.

**Figure 3.**
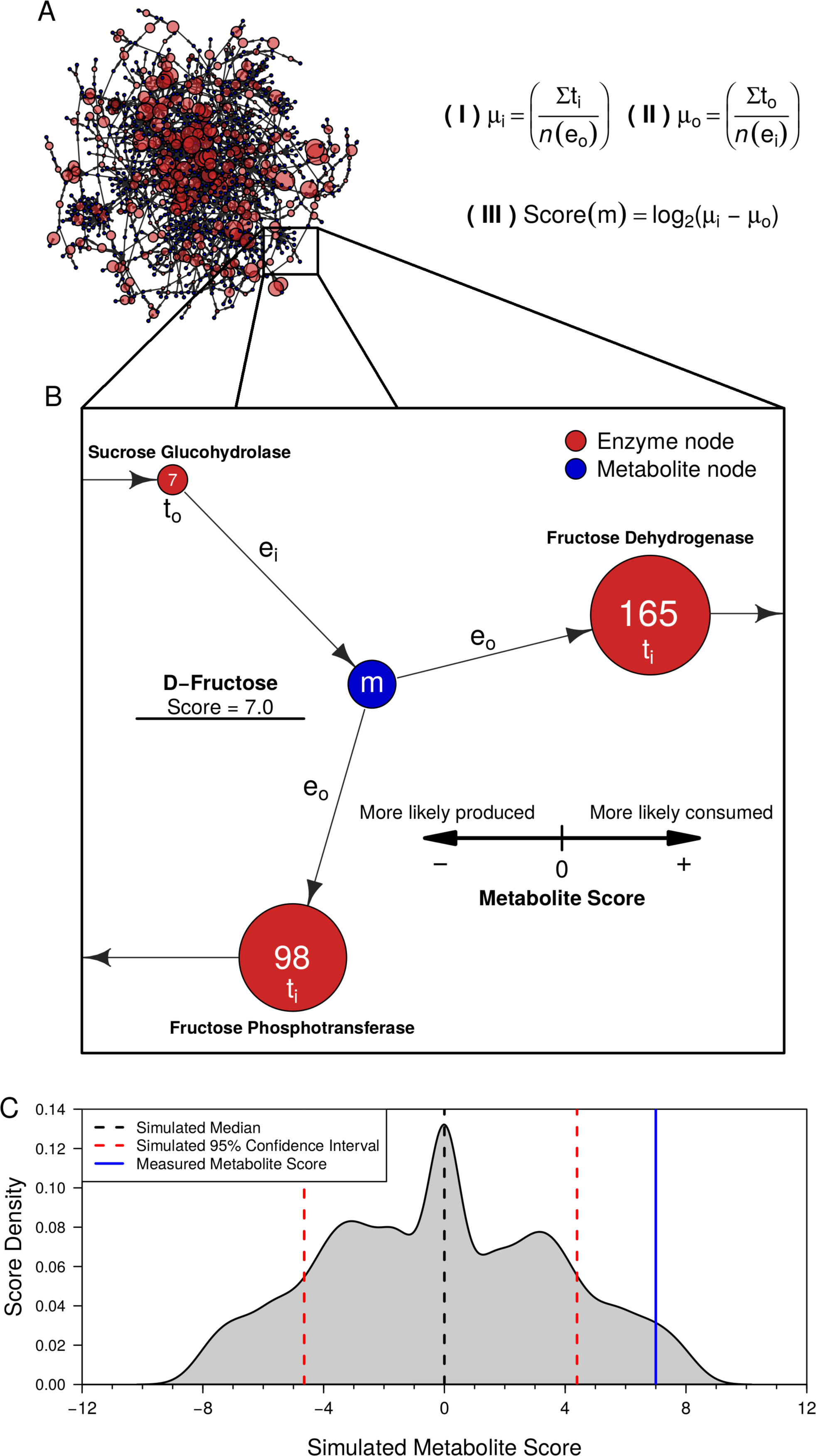
*C. difficile* str. 630 genome-enabled bipartite metabolic network architecture and transcriptomic-enabled metabolite score calculation. **(A)** Largest component from the bipartite GEM of *C. difficile* str. 630. Enzyme node sizes reflect the levels of detectable transcript from each gene. Metabolite score algorithm components: (I) average transcription of reactions consuming a metabolite, (II) average transcription of reactions producing a metabolite, and (III) difference of consumption and production. **(B)** The expanded window displays a partial example of D-fructose score calculation. Values in the red nodes represent normalized transcript reads mapping to enzymes. **(C)** Example 10000-fold Mont-Carlo simulation results corresponding to a significant metabolite score for **m**.

Following application of both methods, we found 5 enzymes that were shared between the top 10 enzymes from BC and CC calculations (2-dehydro-3-deoxyphosphogluconate aldolase, aspartate aminotransferase, pyruvate-flavodoxin oxidoreductase, formate C-acetyltransferase, and 1-deoxy-D-xylulose-5-phosphate synthase). These enzymes primarily participate in core processes including glycolysis, the pentose phosphate pathway, or the citric acid cycle. Upon analysis of the other 15 high-scoring enzymes combined from BC and CC analyses, the majority were also components of the previously mentioned pathways, as well as several for the metabolism of amino acids (Table S3). Similarly, the intersection of those substrates with high both BC and CC values indicated that 6 metabolites were central to the metabolism of *C. difficile* (pyruvate, acetyl-CoA, 2-oxoglutarate, D-4-hydroxy-2-oxoglutarate, D-glyceraldehyde 3-phosphate, and L-glutamate). Not only are these members of glycolysis and the citric acid cycle, but pyruvate, acetyl-CoA, and L-glutamate contribute to numerous intracellular pathways as forms of biological “currency” (39). Notably absent from the most well-connected metabolites were molecules like ATP or NADH. Their exclusion is likely a byproduct of the KEGG LIGAND reference used for network construction, which excludes cofactors from most biochemical reactions. While this may be a limitation of certain analyses, our study was not affected as the primary interest was in those substrates acquired from the environment. These results reflected the defined biological patterns of *C. difficile* and was therefore a viable platform to study metabolism of the pathogen.

**Metabolite score algorithm reveals adaptive nutritional strategies of *C. difficile* during infection of distinct environments.** We next sought to include the transcriptomic results into the metabolic model to infer which metabolites *C. difficile* most likely utilized from a given environment. To accomplish this we mapped normalized transcript abundances to the enzyme nodes in the network. Similar approaches have been previously successful in demonstrating that transcript abundance data can be utilized through the lens of genome-scale metabolic networks to accurately predict microbial metabolic responses to environmental perturbation and identify reporter metabolites of changes (40). In our system, the score of each metabolite was measured as the log_2_-transformed difference in average transcript levels of enzymes that use the metabolite as a substrate and those that generate the metabolite as a product (Fig. 3B). A metabolite with a high score was likely obtained from the environment because the expression of genes for enzymes that produce the metabolite were low. It is important to note that molecules that were more likely produced in our model were not necessarily likely to be released to the environment. Our models do not include the synthesis of large macromolecules (ie. long polypeptides or cytoskeleton) and should therefore only be utilized to consider metabolites that were inputs to a network. Due to the previously mentioned limited technical replication of sequencing efforts, we adopted a Monte Carlo-style simulation for iterative random transcriptome comparison to provide statistical validation of our network-based findings. This process generated random score distributions for each metabolite node in the network, which made it possible to calculate a confidence interval that represented random noise for each metabolite. This ultimately allowed for assessment of the probability that a given metabolite was excluded from the associated null distribution (Fig. 3C).

To identify the metabolites that were most essential for *C. difficile* growth, regardless of the environment, we cross-referenced the 40 highest scoring metabolites from each treatment group (Fig. 4A). N-acetylglucosamine (GlcNAc) was found to the have the highest median score of all shared metabolites, which has been shown to be a readily available source of carbon and nitrogen which can be limiting in the gut (21). We went on to confirm that our strain of *C. difficile* could metabolize GlcNAc for growth (Fig. 4B; Table S5) in *C. difficile* minimal media (41). The Stickland fermentation acceptor proline scored highly across all conditions (42). *C. difficile* is auxotrophic for not only proline, but also cysteine, leucine, isoleucine, tryptophan, and valine, which prevented testing for *in vitro* growth changes on proline despite providing for modest growth in the no carbohydrate control. Previous analysis of *C. difficile* colonizing GF mice under mono-associated conditions indicated that *C. difficile* uses both sets of metabolites (21); however, use of these metabolites in the context of a complex community of potential competitors has not been observed. This analysis indicated that these metabolites might be an integral component of the nutrient niche for *C. difficile*.

**Figure 4.**
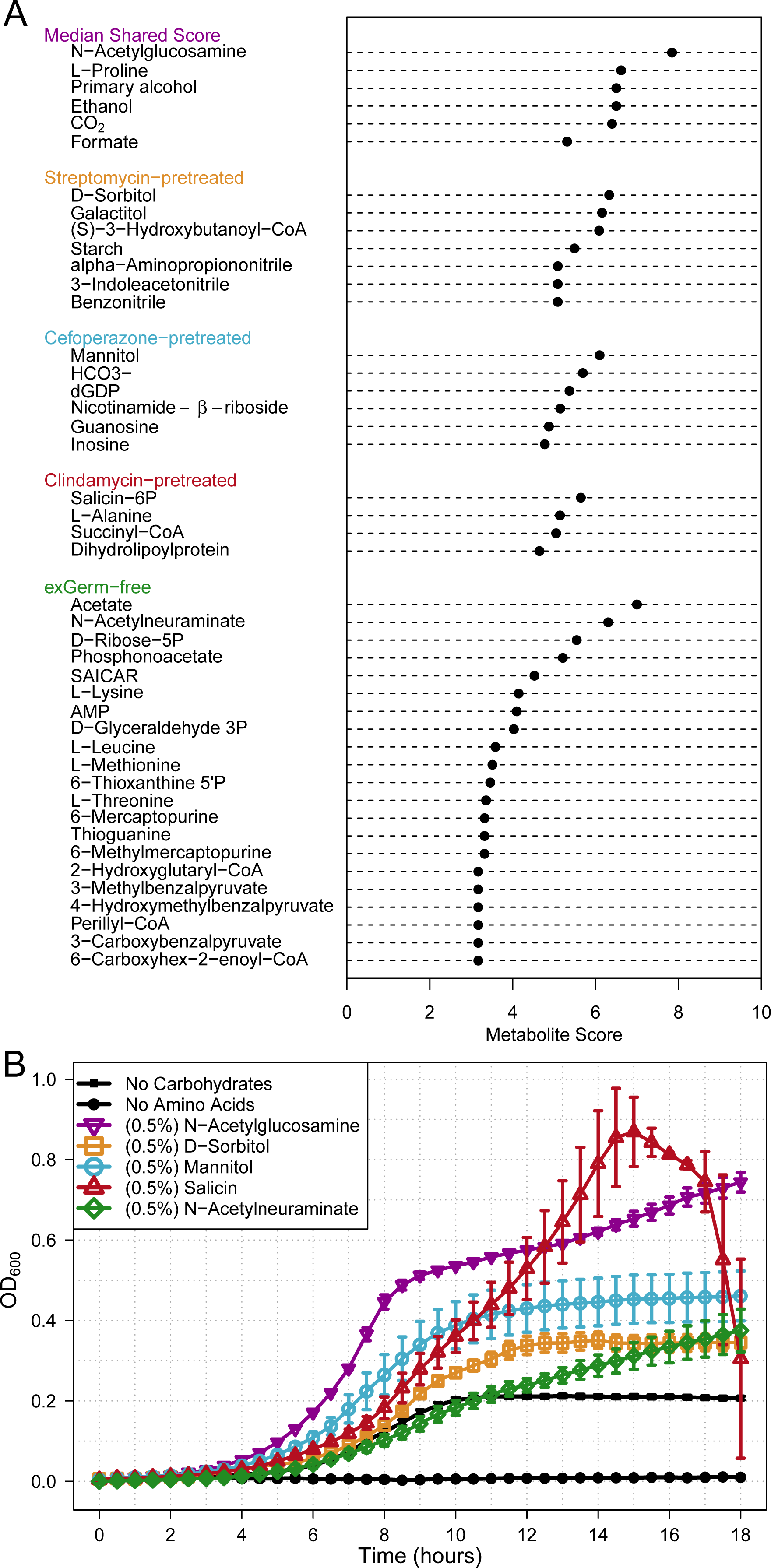
Metabolic network analysis reveals differential carbon source utilization by *C. difficile* across infections. Reported metabolite scores were calculated to have <2.5% probability to be included in the associated random score distribution. Analysis was performed using the 40 highest scoring metabolites from each condition. **(A)** Shared metabolite score represents the median score of metabolites that were consistently scored highly among all infected conditions. Below the conserved patterns, are shown the distinct metabolites for each group’s subset. **(B)** 18 hour *C. difficile* str. 630 *in vitro* growth validating substrates from network analysis. All statistical comparison was performed relative to no carbohydrate control (all *P* < 0.001). Significance was determined with one-way ANOVA with Benjamini–Hochberg correction.

***In vivo* metabolomic analysis supports that *C. difficile* consumes metabolites indicated by metabolic modeling.** To further validate the results of our metabolic model, we tested the effect of *C. difficile* on the metabolite pool in cecal contents from each antibiotic-pretreated and exGF mouse used in the previous analyses. This afforded us the ability to compare replicates within each treatment group. To measure metabolite concentrations, we utilized non-targeted ultra-performance liquid chromatography and mass spectrometry (UPLC-MS) to measure the relative *in vivo* concentrations of metabolites for each mouse in the conditions investigated, with special attention to those highlighted by large metabolite scores. We tested whether the susceptible communities had significantly different concentrations of each metabolite relative to untreated SPF mice and whether the presence of *C. difficile* affected the metabolite composition.

First, we compared the relative concentration of highly scored metabolites in untreated SPF mice and antibiotic pretreated mice in the absence of *C. difficile* (Fig. 5). We found that the relative concentration of GlcNAc was actually significantly lower in all susceptible conditions (Fig. 5A; all *P* < 0.001). The Stickland fermentation acceptors proline (all *P* < 0.05) and hydroxyproline (all *P* < 0.05) were significantly higher in all susceptible environments tested (Fig. 5B and S5D). Succinyl-CoA scored most highly in the clindamycin pretreatment, which is the direct precursor to succinate by succinyl-CoA transferases (43). Succinate has been shown to support *C. difficile* growth *in vivo* through a synergistic relationship that requires at least one other bacterial species (9). As succinyl-CoA was not measured in our metabolimic assay, we instead found that succinate was indeed significantly higher in clindamycin pretreated mice (Fig. 5D; all *P* < 0.05). Among the cefoperazone-pretreated SPF and GF mice, we also found that the concentration of mannitol/sorbitol (Fig. 5C), N-acetylneuraminate (Fig. 5E), and glycine (Fig. S5E) were significantly higher in cefoperazone-treated SPF and GF mice (all *P* < 0.05). These results supported the assertion that susceptible mice had multiple nutrient niches that *C. difficile* was able to exploit.

**Figure 5.**
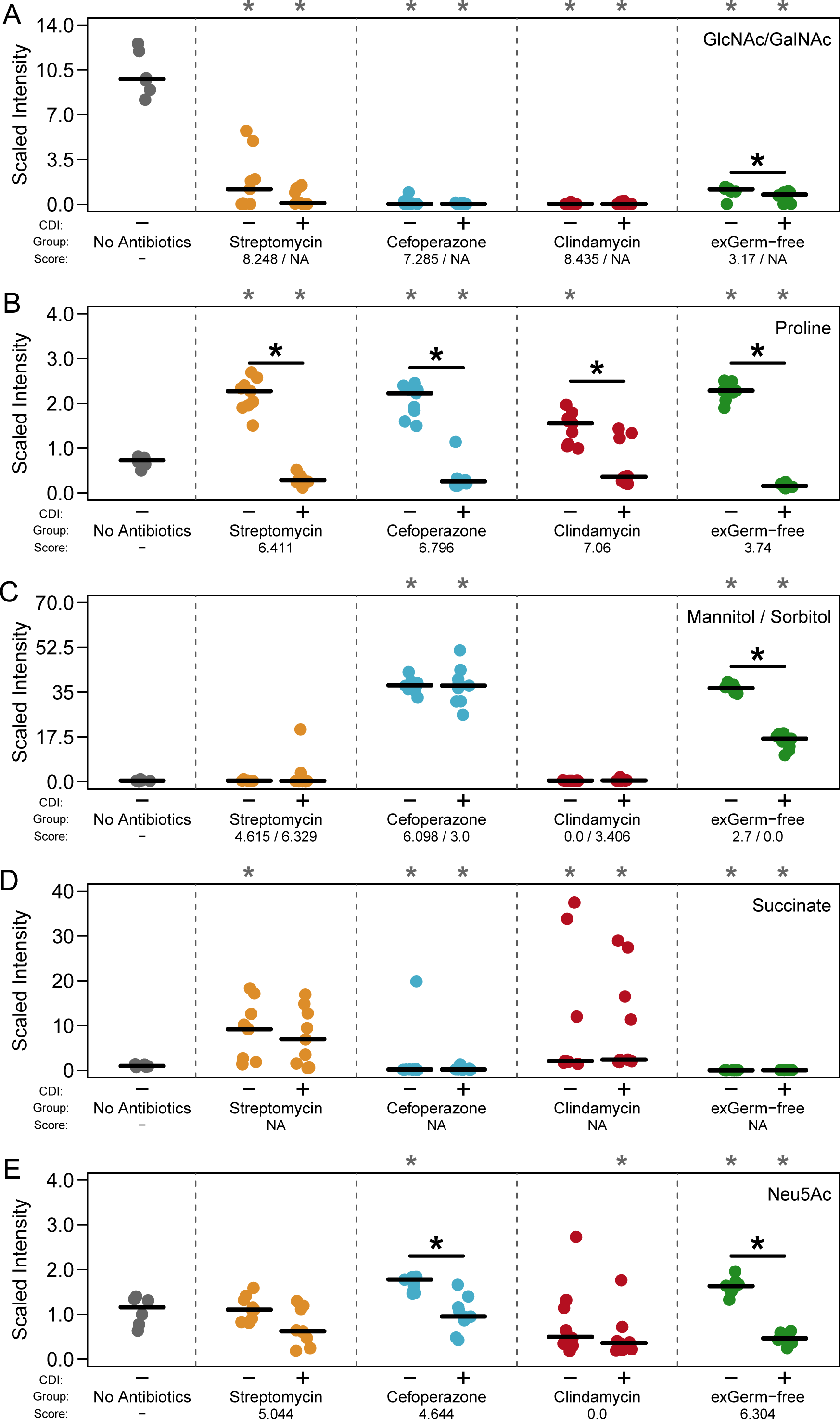
*In vivo* untargeted metabolomics support network-based metabolite scores and suggest nutrient preference hierarchy. Paired metabolites were quantified simultaneously as the only differ by chirality making differentiation impossible. CDI status and *C. difficile* metabolite scores during infection are indicated below each panel. NAs denote metabolites that were not included in our metabolic model of *C. difficile* str. 630. Black asterisks inside the panels represent significant differences between mock and *C. difficile*-infected groups within separate treatment groups (all *P* < 0.05). Gray asterisks along the top margin of each panel indicate significant difference from untreated SPF mice (all *P* < 0.05). Significance was determined with Wilcoxon signed-rank test with Benjamini–Hochberg correction.

Second, we compared relative concentrations of high scoring metabolites during CDI and mock-infection within each pretreatment group (Fig. 5). Both groups of host-derived glycans, GlcNAc/GalNAc (Fig. 5A) and Neu5Ac (Fig. 5E), were significantly lower when in the presence of *C. difficile* in exGF mice (*P* < 0.05 and 0.01). In agreement with the previous results, we found that the Stickland acceptors proline (Fig. 5B) and hydroxyproline (Fig. S5D) were significantly lower in every *C. difficile* colonized environment (all *P* < 0.05). Glycine, another preferred Stickland acceptor, was lower in each condition following infection with significant change in cefoperazone-pretreated mice (Fig. S5D; *P* < 0.05). The Stickland donors leucine and isoleucine were significantly lower in all infected conditions except streptomycin-pretreated mice (Fig. S5; all *P* < 0.05). These results supported the hypothesis that amino acids are an important energy source of *C. difficile* during infection. A significant difference was seen for mannitol/sorbitol in exGF mice (*P* < 0.01), but not in cefoperazone-pretreated mice (Fig. 5C). Although a lower concentration of succinate in both streptomycin and clindamycin pretreated mice was observed, neither was found to be significant. Overall, this metabolomic analysis supported our transcriptome-based metabolite score algorithm for predicting the metabolites utilized by *C. difficile* during different infection conditions. Results from metabolic modeling combined with untargeted metabolomic analysis also suggested a possible hierarchy of preferred growth substrates.

## Discussion

The results presented here expand upon previous understanding of *C. difficile* metabolism during infection by showing that not only does the pathogen adapt its metabolism to life inside of a host (14, 21), but also to the context of the specific gut environment in which it finds itself. Previous transcriptomic efforts to measure the response of *C. difficile* have characterized *in vivo* changes in metabolism following colonization of GF mice. In this study, we utilized a conventionally-reared mouse model of infection to compare the response of *C. difficile* to colonization in the context of varied gut communities generated by pretreatment with representatives from distinct classes of antibiotics. With these models, we identified subtle differences in sporulation and toxin activity between each antibiotic-pretreated condition. Transcriptomic sequencing of *C. difficile* across colonized environments indicated complex expression patterns of genes in catabolic pathways for a variety of carbon sources. Through integration of transcriptomic data with genome-scale metabolic modeling allowed us to observe that *C. difficile* likely generated energy by metabolizing specific alternative carbon and nitrogen sources across colonized conditions. We also found that Stickland fermentation substrates and products, as well as the host-derived glycan N-acetylglucosamine, were consistently among the highest scoring shared metabolites which indicated that these metabolites were central to the *in vivo* nutritional strategy of *C. difficile*. To confirm our modeling-based results we employed untargeted mass spectrometry that demonstrated greater availability of many metabolites highlighted by our algorithm in susceptible gut environments. Metabolomic analysis further revealed differential reduction of highly scored metabolites during CDI, which suggested a hierarchy for the utilization of certain growth nutrients.

Our interpretation of the positive trends we observed between metabolite score and substrate availability across conditions was that the distinct antibiotic treatments eliminate alternative patterns of competitors for those nutrients in the gut of susceptible animals. These groups of bacteria are likely to be more specialized than *C. difficile* at acquiring those resources, supporting the nutrient-niche hypothesis as a mechanism to explain the exclusion of *C. difficile* by the intact microbiota. By pursuing a more generalist behavior in terms of growth nutrient preferences, *C. difficile* has increased fitness for exploiting differentially perturbed gut communities. ExGF mice provided a partially controlled system of resource competition. In this condition, Neu5Ac was found to be the highest scored substrate and its concentrations were significantly higher during mock infection than following *C. difficile* colonization. A similar trend was also seen in cefoperazone-pretreated mice, implying that this antibiotic may have reduced the population density of the particular competitors for this niche. These data suggest that *C. difficile* may be less competitive for this host-derived glycan and only has access when certain competitors have had their densities reduced. In agreement with earlier research we found that *C. difficile* likely fermented amino acids for energy during infection of GF mice in addition to host-derived glycan catabolism. Our results go on to support that this metabolic strategy was conserved across all infection conditions tested. Several Stickland substrates had consistently high metabolite scores including alanine, leucine, and proline indeed dropped concentration during infection (Table S4, Fig. 5B, S5A, and S5B). Fermentation of amino acids provides not only carbon and energy, but are also a source of nitrogen which is a limited resource in the mammalian lower gastrointestinal tract (44). This makes Stickland fermentation a valuable metabolic strategy, and it stands to reason that *C. difficile* would use this strategy across all environments it colonizes. This same principle may also extend to glycans harvested from the host mucus layer (GlcNAc and Neu5Ac) as they are another source of carbon and nitrogen which, despite augmented release by members of the microbiota, would be present at some basal concentration regardless of other species’ metabolism (45, 46). Moreover, decreases in relative concentration of certain metabolites following antibiotic treatment does not preclude their availability to *C. difficile*. As long as competition for the remaining pool of the given substrate is reduced, *C. difficile* may be able to exploit it as a component of its nutrient niche space. Based on our results, we propose that amino acid catabolism is a primary strategy of *C. difficile in vivo* followed closely by host-derived glycans catabolism. To fulfill its remaining needs, *C. difficile* adapts its metabolism to utilize a combination of carbohydrates, sugar alcohols, or carboxylic acids depending on their availability in the environment. Since the latter provide carbon and energy but not nitrogen, it appears that *C. difficile* metabolism strongly prefers nitrogen-containing carbon sources that fulfill a larger proportion of its biological requirements.

Several factors limited our ability to generate transcriptomic replicates for individual mice in each treatment group. Most prominently, we were forced to pool the cecal contents of multiple animals to generate a sufficient quantity of high quality RNA and sequence extremely deeply that would permit sampling the transcriptome of a rare member of the microbiota (Fig. S2C). Due to possible variation between individual samples that could be masked by this approach, we quantified within-group sample variation for all sample types for which we were able to collect biological replicates. This included *C. difficile* density, 16S rRNA gene abundance, and untargeted mass-spectrometry. In order to increase our confidence that transcriptomes were more likely to be consistent between pretreatment groups, we calculated within-group sample variance for *C. difficile* density, 16S rRNA gene abundances, and untargeted metabolomics datasets (Fig. S3B-D). This revealed extremely low variability in each treatment group tested for sample types with increasing levels of complexity. Since these data were collected using matched cecal samples, we were confident that our transcriptomic results reflected reality. Unlike our transcriptomic data, we were able to characterize the metabolome of individual animals; however, these comparisons had their own complications related to the fact that multiple organisms contribute to the overall metabolite pool. The changes observed could be the result of metabolic patterns from other species in each system (host or microbe) in response to pathogen colonization, and it is difficult to discern whether *C. difficile* reaches a density large enough to impact these differences on its own. Possible limitations of our modeling approach also existed, despite much of our results being consistent with previously published work and our own untargeted metabolomic analysis. The metabolite score calculation is dependent on correct and existing gene annotation. In this regard it has been shown that the pathway annotations in KEGG are robust to missing elements (47), however this does not completely eliminate the possibility for this type of error. Due to the topology of the metabolic network, we were also unable to integrate stoichiometry for each reaction which may effect rates of consumption or production. Reaction reversibility also varies depending on versions of enzymes possessed by each species. Since our algorithm favorably weights those metabolites closer to the network periphery, incorrect directionality annotations may lead to mislabeling reactants or products and potentially lead to incorrect metabolite score calculations. Since our metabolite scoring algorithm selectively amplifies signal for those metabolites with the highest probability to be imported from the environment, this modeling platform may also allow for the identification of emergent properties for the metabolism of *C. difficile* during infection. One example could be the appearance of CO_2_ and formate, apparent metabolic end products, in the list of shared metabolites which scored highly across conditions. Although this may be a shortcoming of the genome or database annotation, one group has posited that *C. difficile* may actually consume CO_2_ under certain conditions and require both of these substrates to undergo this process (48). These findings highlight that our method not only identified growth substrates, but also identified additional metabolites that were being utilized for other processes. With further manual curation of the *C. difficile* metabolic network, more species-specific discoveries are possible. Even with this possibility, the application of multiple methods to study the altered physiology of *C. difficile* in mock-infected and infected communities allowed us to validate our results based on known elements of *C. difficile* biology and to internally cross validate the novel results from our experiments. Ultimately, these results combine to underscore predictions of nutrient niche plasticity.

Our systems approach to studying *C. difficile* metabolism during the infection of susceptible communities combines multiple levels of biological data to identify metabolic trends that would not be apparent by a single method. Only through integrative multi-omic analysis of *C. difficile* infection employing genomics, transcriptomics, and metabolomics were we able to uncover a much clearer image of *C. difficile*’s nutrient niche space during infection in the context of complex microbial communities. Focusing on previously established metabolic capabilities of the pathogen, we identify that these forms of metabolism are differentially important to *C. difficile* when colonizing distinct environments. Our data suggest that *C. difficile* is true bacterial generalist, making it less competitive for specific nutrients against specialists, but more fit overall for colonizing a variety of recently vacated nutrient niche spaces. These results have implications for the development of targeted measures to prevent *C. difficile* colonization through pre- or probiotic therapy that will need to be tailored to specific antibiotic-induced perturbations. In the future, this systems-level approach could be expanded to study the niche landscape of entire communities of bacteria and subsequent changes to competition for nutrients in response to antibiotic treatment or pathogen colonization.

## Materials and Methods

**Animal care and antibiotic administration.** Six-to-eight week-old GF C57BL/6 mice were obtained from a single breeding colony maintained at the University of Michigan and fed Laboratory Rodent Diet 5001 from LabDiet for all experiments. All animal protocols were approved by the University Committee on Use and Care of Animals at the University of Michigan and carried out in accordance with the approved guidelines. Specified SPF animals were administered one of three antibiotics; cefoperazone, streptomycin, or clindamycin (Table 1). Cefoperazone (0.5 mg/ml) and streptomycin (5.0 mg/ml) were administered in distilled drinking water *ad libitum* for 5 days with 2 days recovery with untreated distilled drinking water prior to infection. Clindamycin (10 mg/kg) was given via intraperitoneal injection 24 hours before time of infection. Adapted from a previously described model (24).

***C. difficile* infection and necropsy.** All *C. difficile* strain 630 spores were prepared from a single large batch whose concentration was determined a week prior to challenge. On the day of challenge, 1×10^3^ *C. difficile* spores were administered to mice via oral gavage in phosphate-buffered saline (PBS) vehicle. Subsequent quantitative plating to enumerate the spores was performed to ensure correct dosage. Mock-infected animals were given an oral gavage of 100 μl PBS at the same time as those mice administered *C. difficile* spores. 18 hours following infection, mice were euthanized by carbon dioxide asphyxiation and necropsied to obtain the cecal contents. Two 100 μl aliquots were immediately flash frozen for later DNA extraction and toxin titer analysis, respectively. A third 100 μl aliquot was quickly transferred to an anaerobic chamber for quantification of *C. difficile* abundance. The remaining content in the ceca (approximately 1 mL) was mixed with 1 mL of sterile PBS in a stainless steel mortar housed in a dry ice and ethanol bath. The cecal contents of 9 mice from 3 cages was pooled into the mortar. Pooling cecal contents was necessary so that there would be a sufficient quantity of high quality rRNA-free RNA for deep sequencing. The pooled content was then finely ground and stored at -80° C for subsequent RNA extraction.

***C. difficile* cultivation and quantification.** Cecal samples were weighed and serially diluted under anaerobic conditions (6% H, 20% CO_2_, 74% N_2_) with anaerobic PBS. Differential plating was performed to quantify both *C. difficile* spores and vegetative cells by plating diluted samples on CCFAE plates (fructose agar plus cycloserine (0.5%), cefoxitin (0.5%), and erythromycin (0.2%)) at 37° C for 24 hours under anaerobic conditions (49). It is important to note that the germination agent taurocholate was omitted from these plates to quantify only vegetative cells. In parallel, undiluted samples were heated at 60° C for 30 minutes to eliminate vegetative cells and leave only spores (50). These samples were serially diluted under anaerobic conditions in anaerobic PBS and plated on CCFAE with taurocholate (10%) at 37° C for 24 hours. Plating was simultaneously done for heated samples on CCFAE to ensure all vegetative cells had been eliminated.

***C. difficile* toxin titer assay.** To quantify the titer of toxin in the cecum a Vero cell rounding assay was performed as in (27). Briefly, filtered-sterilized cecal content was serially diluted in PBS and added to Vero cells in a 96-well plate. Plates were blinded and viewed after 24 hour incubation for cell rounding. A more detailed protocol with product information can be found at: https://github.com/SchlossLab/Jenior_Modeling_mSystems_2017/blob/master/protocols/toxin_assay/Verocell_ToxinActivity_Assay.Rmd

**16S rRNA gene sequencing and read curation.** DNA was extracted from approximately 50 mg of cecal content from each mouse using the PowerSoil-htp 96 Well Soil DNA isolation kit (MO BIO Laboratories) and an epMotion 5075 automated pipetting system (Eppendorf). The V4 region of the bacterial 16S rRNA gene was amplified using custom barcoded primers and sequenced as described previously using an Illumina MiSeq sequencer (51). All 63 samples were sequenced on a single sequencing run. The 16S rRNA gene sequences were curated using the mothur software package (v1.36), as described previously (51). In short, paired-end reads were merged into contigs, screened for quality, aligned to SILVA 16S rRNA sequence database, and screened for chimeras. Sequences were classified using a naive Bayesian classifier trained against a 16S rRNA gene training set provided by the Ribosomal Database Project (RDP) (52). Curated sequences were clustered into operational taxonomic units (OTUs) using a 97% similarity cutoff with the average neighbor clustering algorithm. The number of sequences in each sample was rarefied to 2,500 per sample to minimize the effects of uneven sampling.

**RNA extraction, shotgun library preparation, and sequencing.** Pooled, flash-frozen samples were ground with a sterile pestle to a fine powder and scraped into a sterile 50 ml polypropylene conical tube. Samples were stored at -80° C until the time of extraction. Immediately before RNA extraction, 3 ml of lysis buffer (2% SDS, 16 mM EDTA and 200 mM NaCl) contained in a 50 ml polypropylene conical tube was first heated for 5 minutes in a boiling water bath (53). The hot lysis buffer was added to the frozen and ground cecal content. The mixture was boiled with periodic vortexing for another 5 minutes. After boiling, an equal volume of 37° C acid phenol/chloroform was added to the cecal content lysate and incubated at 37° C for 10 minutes with periodic vortexing. The mixture was the centrifuged at 2,500 x g at 4° C for 15 minutes. The aqueous phase was then transferred to a sterile tube and an equal volume of acid phenol/chloroform was added. This mixture was vortexed and centrifuged at 2,500 x g at 4° for 5 minutes. The process was repeated until aqueous phase was clear. The last extraction was performed with chloroform/isoamyl alcohol to remove the acid phenol. An equal volume of isopropanol was added and the extracted nucleic acid was incubated overnight at -20° C. The following day the sample was centrifuged at 12000 x g at 4° C for 45 minutes. The pellet was washed with 0° C 100% ethanol and resuspended in 200 μl of RNase-free water. Samples were then treated with 2 μl of Turbo DNase for 30 minutes at 37° C. RNA samples were retrieved using the Zymo Quick-RNA MiniPrep. Completion of the DNase reaction was assessed using PCR for the V4 region of the 16S rRNA gene for 30 cycles (Kozich, 2013). Quality and integrity of RNA was measured using the Agilent RNA 6000 Nano kit for total prokaryotic RNA. The Ribo-Zero Gold rRNA Removal Kit Epidemiology was then used to deplete 16S and 18S rRNA from the samples. Prior to library construction, quality and integrity as measured again using the Agilent RNA 6000 Pico Kit. Stranded RNA-Seq libraries were made constructed with the TruSeq Total RNA Library Preparation Kit v2. The Agilent DNA High Sensitivity Kit was used to measure concentration and fragment size distribution before sequencing. High-throughput sequencing was performed by the University of Michigan Sequencing Core in Ann Arbor, MI. For all groups, sequencing was repeated across 4 lanes of an Illumina HiSeq 2500 using the 2x50 bp chemistry.

**cDNA read curation, mapping, and normalization.** Raw read curation was performed in a two step process. First, residual 5’ and 3’ Illumina adapter sequences were removed using CutAdapt (54) on a per library basis. Reads were then quality trimmed using Sickle (Joshi, 2011) on the default settings. An average of ∼261,000,000 total reads (both paired and orphaned) remained after quality trimming. Mapping was accomplished using Bowtie2 (55) and the default stringent settings allowing for 0 mismatches again target reference genes. An average of ∼6,880,000 reads in sample each mapped to the annotated nucleotide gene sequences of *Clostridioides difficile* 630 from the KEGG: Kyoto Encyclopedia of Genes and Genomes (56). Optical and PCR duplicates were then removed using Picard MarkDuplicates (http://broadinstitute.github.io/picard/), leaving an average of ∼167,000 reads per sample for final analysis (Table S2). The remaining mappings were converted to idxstats format using Samtools (57) and the read counts per gene were tabulated. Discordant pair mappings were discarded and counts were then normalized to read length and gene length to give a per base report of gene coverage. Each collection of reads was then subsampled to 90% of the lowest sequence total across the libraries resulting in even quantities of normalized read abundances in each group to be utilized in downstream analysis. This method was chosen as normalization to housekeeping genes would artificially remove their contributions to metabolic flux and reduce the information provided by our metabolite score calculations within our metabolic modeling approach.

**Reaction Annotation & Bipartite Network Construction.** The metabolism of *C. difficile* stain 630 was represented as a directed bipartite graph with both enzymes and metabolites as nodes. Briefly, models were semi-automatically constructed using KEGG (2016 edition) ortholog (KO) gene annotations to which transcripts had been mapped. Reactions that each KEGG ortholog mediate were extracted from ko_reaction.list located in /kegg/genes/ko/. KOs that do not mediate simple biochemical reactions (e.g. mediate interactions of macromolecules) were omitted. Metabolites linked to each reaction were retrieved from reaction_mapformula.lst file located in /kegg/ligand/reaction/ from the KEGG release. Those reactions that did not have annotations for the chemical compounds the interact with are discarded. Metabolites were then associated with each enzyme and the directionality and reversibility of each biochemical conversion was also saved. This process was repeated for all enzymes in the given bacterial genome, with each enzyme and metabolite node only appearing once. The resulting data structure was an associative array of enzymes associated with lists of both categories of substrates (input and output), which could then be represented as a bipartite network. The final metabolic network of C. difficile strain 630 contained a total of 1205 individual nodes (447 enzymes and 758 substrates) with 2135 directed edges. Transcriptomic mapping data was then re-associated with the respective enzyme nodes prior to scoring calculations. Betweenness-centrality and overall closeness centralization indices were calculated using the igraph R package found at http://igraph.org/r/.

**Metabolite Score Calculation.** The substrate scoring algorithm (Fig. 3A) favors metabolites that are more likely acquired from the environment (not produced within the network), and will award them a higher score (Fig. 3B & 4A). The presumption of our approach was that enzymes that were more highly transcribed were more likely to utilize the substrates they act on due to coupled bacterial transcription and translation. If a compound was more likely to be produced, the more negative the resulting score would be. To calculate the score of a given metabolite (m), we used rarefied transcript abundances mapped to respective enzyme nodes. This was represented by t_o_ and t_i_ to designate if an enzyme created or utilized m. The first step was to calculate the average expression of enzymes for reactions that either created a given metabolite (i) or consumed that metabolite (ii). For each direction, the sum of transcripts for enzymes connecting to a metabolite were divided by the number of contributing edges (e_o_ or e_i_) to normalize for highly connected metabolite nodes. Next the raw metabolite score was calculated by subtracting the creation value from the consumption value to weight for metabolites that are likely acquired exogenously. The difference was log_2_ transformed for comparability between scores of individual metabolites. This resulted in a final value that reflected the likelihood a metabolite was acquired from the environment. Untransformed scores that already equaled to 0 were ignored and negative values were accounted for by transformation of the absolute value then multiplied by -1. These methods have been written into a single python workflow, along with supporting reference files, and is presented as bigSMALL v1.0 (BacterIal Genome-Scale Metabolic models for AppLied reverse ecoLogy) available in a public Github repository at https://github.com/mjenior/bigsmall.

**Transcriptome Randomization and Probability Distribution Comparison.** As sequencing replicates of *in vivo* transcriptomes was not feasible, we applied a Monte Carlo style simulation to distinguish calculated metabolite scores due to distinct transcriptional patterns for the environment measured from those metabolites that were constitutively scored at the extremes of the scale. We employed a 10,000-fold bootstrapping approach of randomly reassigning transcript abundance for enzyme nodes and recalculating metabolite scores. This approach was chosen over fitting a simulated transcriptome to a negative binomial distribution because it created a more relevant standard of comparison for lower coverage sequencing efforts. Using this method, each substrate node accumulated a random probability distribution of metabolite scores which were then used to calculate the median and confidence interval to generate a probability for each metabolite score to be the result of more than chance. This was a superior approach to switch randomization since the connections of the network itself was created through natural selection and any large-scale alterations would yield biologically uninformative comparisons (58).

**Anaerobic *in vitro C. difficile* growth curves.** The carbon-free variation of *C. difficile* Basal Defined Medium (NCMM) was prepared as previously described (6). Individual carbohydrate sources were added at a final concentration of 5 mg/mL and pair-wise carbohydrate combinations were added at 2.5 mg/mL each (5 mg/mL total). A solution of the required amino acids was made separately and added when noted at identical concentrations to the same study. 245 μl of final media mixes were added to a 96-well sterile clear-bottom plate. A rich media growth control was also included, consisting of liquid Brain-Heart Infusion with 0.5% cysteine. All culturing and growth measurement were performed anaerobically in a Coy Type B Vinyl Anaerobic Chamber (3.0% H, 5.0% CO_2_, 92.0% N, 0.0% O_2_). *C. difficile* str. 630 was grown for 14 hours at 37° C in 3 mL BHI with 0.5% cysteine. Cultures were then centrifuged at 2000 rpm for 5 minutes and resulting pellets were washed twice with sterile, anaerobic phosphate-buffered saline (PBS). Washed pellets were resuspended in 3 mL more PBS and 5 μl of prepped culture was added the each growth well of the plate containing media. The plate was then placed in a Tecan Sunrise plate reader. Plates were incubated for 24 hours at 37° C with automatic optical density readings at 600 nm taken every 30 minutes. OD_600_ values were normalized to readings from wells containing sterile media of the same type at equal time of incubation. Growth rates and other curve metrics were determined by differentiation analysis of the measured OD_600_ over time in R to obtain the slope at each time point.

**Quantification of *in vivo* metabolite relative concentrations.** Metabolomic analysis performed by Metabolon (Durham, NC), a brief description of their methods is as follows. All methods utilized a Waters ACQUITY ultra-performance liquid chromatography (UPLC) and a Thermo Scientific Q-Exactive high resolution/accurate mass spectrometer interfaced with a heated electrospray ionization (HESI-II) source and Orbitrap mass analyzer at 35,000 mass resolution. Samples were dried then reconstituted in solvents compatible to each of the four methods. The first, in acidic positive conditions using a C18 column (Waters UPLC BEH C18-2.1x100 mm, 1.7 µm) using water and methanol, containing 0.05% perfluoropentanoic acid (PFPA) and 0.1% formic acid (FA). The second method was identical to the first but was chromatographically optimized for more hydrophobic compounds. The third approach utilized a basic negative ion optimized conditions using a separate dedicated C18 column. Basic extracts were gradient eluted from the column using methanol and water, however with 6.5mM Ammonium Bicarbonate at pH 8. Samples were then analyzed via negative ionization following elution from a hydrophilic interaction chromatography column (Waters UPLC BEH Amide 2.1x150 mm, 1.7 µm) using a gradient consisting of water and acetonitrile with 10mM Ammonium Formate, pH 10.8. The MS analysis alternated between MS and data-dependent MS n scans using dynamic exclusion. The scan range varied slighted between methods but covered 70-1000 m/z. Library matches for each compound were checked for each sample and corrected if necessary. Peaks were quantified using area under the curve.

**Statistical methods.** All statistical analyses were performed using R (v.3.2.0). Significant differences between community structure of treatment groups from 16S rRNA gene sequencing were determined with AMOVA in the mothur software package. Significant differences of Inv. Simpson diversity, cfu, toxin titer, and metabolite concentrations were determined by Wilcoxon signed-rank test with Benjamini-Hochberg correction. Undetectable points used half the limit of detection for all statistical calculations. Significant differences for growth curves compared to no carbohydrate control (+ amino acids) were calculated using 1-way ANOVA with Benjamini-Hochberg correction.

## Acknowledgements

The authors would like to acknowledge Charles Koumpouras for assistance with DNA extractions and metabolomic sample preparation. We would also like to acknowledge members of the University of Michigan Germ-free Mouse Center, University of Michigan Sequencing Core, and Metabolon for their assistance in experimental design, execution, and data collection. Pooled and quality trimmed transcriptomic read data and experiment metadata are available through the NCBI Sequence Read Archive (SRA; PRJNA354635). Data processing steps for beginning from raw sequence data to the final manuscript are hosted at http://www.github.com/SchlossLab/Jenior_Modeling_mSystems_2017. The authors would additionally like to thank Geoffrey Hannigan Ph.D, Kaitlin Flynn Ph.D, and Nielsen Baxter Ph.D. for their suggestions on manuscript drafts.

**Author Affiliations Department of Microbiology and Immunology, University of Michigan, Ann Arbor, Michigan.** Matthew L. Jenior, Jhansi L. Leslie, & Patrick D. Schloss Ph.D.

**Department of Internal Medicine/Infectious Diseases Division, University of Michigan Medical Center, Ann Arbor, Michigan. Department of Microbiology and Immunology, University of Michigan, Ann Arbor, Michigan.** Vincent B. Young M.D. Ph.D.

**Author Contributions** M.L.J. conceived, designed and performed experiments, analyzed data, and drafted the manuscript. J.L.L. performed experiments, analyzed data, and contributed to the manuscript. V.B.Y. contributed to the manuscript. P.D.S. interpreted data and contributed the manuscript. The authors declare no conflicts of interest.

**Corresponding author** Correspondence to <u>Patrick D. Schloss</u>

---

**Supplementary Figure 1 Experimental timelines for mouse model pretreatments and *C. difficile* infection.** 9 wild-type C57BL/6 mice across 3 cages were included in each treatment group. **(A)** Streptomycin or **(B)** cefoperazone administered *ad libitum* in drinking water for 5 days with 2 days recovery with untreated drinking water before infection, **(C)** a single clindamycin intraperitoneal injection one day prior to infection, or **(D)** no antibiotic pretreatment (for both SPF control and GF mice). If no antibiotics were administered in the drinking water, mice were given untreated drinking water for the duration of the experiment beginning 7 days prior to infection. At the time of infection, mice were challenged with 1×10^3^ *C. difficile* str. 630 spores. Euthanization and necropsy was done 18 hours post-challenge and cecal content was then collected.

**Supplementary Figure 2 Analysis of bacterial community structure resulting from antibiotic treatment.** Results from 16S rRNA gene amplicon sequencing from bacterial communities of cecal content in both mock-infected and *C. difficile* 630-infected animals 18 hours post-infection across pretreatment models. **(A)** Non-metric multidimensional scaling (NMDS) ordination based on Theta_YC_ distances for the gut microbiome of all SPF mice used in these experiments (n = 36). All treatment groups are significantly different from each other groups by AMOVA (*P* < 0.001). **(B)** Inverse Simpson diversity for each cecal community from the mice in (A). Cecal communities from mice not treated with any antibiotics are significantly more diverse than any antibiotic-pretreated condition (*P* < 0.001). **(C)** Representation of 16S amplicon reads contributed by *C. difficile* in each sequenced condition compared to the total bacterial community. The percents listed at the top of each group is the proportion of the total community represented by *C. difficile*. Significantly less were for *C. difficile* were detected in each condition (*P* < 0.001).

**Supplementary Figure 3 Levels of within-group variation across datasets generated for this study. (A)** Normalized transcript abundance of select housekeeping and central metabolism genes. (I) Housekeeping genes; DNA gyrase subunit A (GyrA), threonyl-tRNA synthetase (ThrS), and ATP-dependent Clp protease (ClpP).(II) Genes in separate metabolic pathways that contribute to input substrate score; enolase, glycine reductase (GrdA), and D-proline reductase (PrdA). **(B)** Median sample variance for vegetative *C. difficile* cfu from each colonized condition. **(C)** Median and interquartile range of the sample variance of OTU abundances from 16S rRNA gene sequencing, sample variances for each OTU were calculated individually prior to summary statistic calculations. **(D)** Median and interquartile range of the sample variance of Scaled intensities from untargeted metabolomic analysis, sample variances for each metabolite were in the same fashion as with OTU abundances. Data (other than transcriptomic results) was collected from the same nine animals per group were (n = 9).

**Supplementary Figure 4 Select *C. difficile* gene set expression compared between treatment group.** Relative abundances of *C. difficile* transcript for specific genes of interest. **(A)** Transcription for select genes from the *C. difficile* sporulation pathway with the greatest variation in expression between the conditions tested. **(B)** Relative abundances of transcript for genes that encode effector proteins from the *C. difficile* pathogenicity locus. **(C)** Transcript abundances for genes associated with quorum sensing in *C. difficile*. **(D)** Transcript relative abundance of select sigma factors which expression or activity is influenced by environmental metabolite concentrations. Asterisks (*) indicate genes from which transcript was undetectable.

**Supplementary Figure 5 Change in *in vivo* concentrations of additional Stickland fermentation substrates.** Comparison of concentrations for other Stickland fermentation substrates from *C. difficile*-infected and mock-infected mouse cecal content 18 hours post-infection. Labels in the top left corner of each panel indicate whether the amino acid is a Stickland donor or acceptor. Black asterisks inside the panels denote significant differences between mock and *C. difficile*-infected groups within separate treatment groups (all *P* < 0.05). Gray asterisks along the top margin of each panel indicate significant difference from untreated SPF mice (all *P* < 0.05).

**Supplementary Table 1 Specific genes and normalized cDNA read abundances included in analysis reported in Figure 2.** Transcript abundances reported in each of the antibiotic associated columns were first normalized to both sequencing read length and target gene length. Each of the three groups were then even subsampled to an equal total sequences abundance of 13,000 reads to allow for comparability between groups. Additional columns indicate specific gene annotation (gene, pathways, & KEDD_ID) as well as which group each gene belongs for ternary plot (family).

**Supplementary Table 2 Normalized cDNA read abundances, gene annotations, and enzymatic reaction information used for metabolic model building for *C. difficile* str. 630 KEGG orthologs across colonized conditions.** All KEGG orthologs included in the *C. difficile* str. 630 KEGG genome annotation (2015) were included in this analysis. Read abundances were normalized as previously outlined to sequencing read length, target gene length, and even total sampling between groups. Also included are individual enzyme annotation for each KEGG ortholog, as well as the associated biochemical reaction information extracted from reaction/reaction_mapformula.lst from KEGG. Together, KEGG ortholog and enzymatic reaction data were used to reconstruct the metabolic network of *C. difficile* str. 630 in presented analyses.

**Supplementary Table 3 Topology metrics for enzyme and metabolite nodes in the *C. difficile* str. 630 metabolic network.** Topology analysis of the metabolic network assembled for this study was performed in the absence of transcriptomic data to assess quality of *de novo* assembled network in its reflection of known bacterial metabolism patterns. Enzyme and metabolite node analysis are presented on separate tabs. Centrality metrics and brief explanations are as follows: Degree is the total number of connections for a given node (both incoming and outgoing), Betweenness is the number of shortest paths connecting all other nodes pairs that pass through the node of interest, and Closeness is the inverse sum of shortest path length that pass through the node of interest. Combined these calculation inform how strongly connected a node is and how vital it is too overall network structure.

**Supplementary Table 4 Metabolites with significant metabolite scores for *C. difficile* in each colonized condition.** Each tab represents those metabolites found to exceed the significance cutoffs for *C. difficile* str. 630 after colonization of each of the respective susceptible states. These threshold were set for each metabolite independently through Monte Carlo simulation as outlined by Figure 3C. A *p*-value of < 0.05 corresponded to a metabolite scoring outside of the 95% confidence interval in the random distribution, and *p* < 0.01 corresponds to those outside the 99% confidence interval. Confidence interval calculations for non-normal distributions were performed as defined by (59).

**Supplementary Table 5 *In vitro* growth analysis for *C. difficile* 630 with carbon sources identified by metabolic network algorithm.** Analysis of growth on highly scored carbon sources to identify possible differences in utilization efficiency.

